# The impact of phage and phage resistance on microbial community dynamics

**DOI:** 10.1101/2023.09.26.559468

**Authors:** Ellinor O Alseth, Rafael Custodio, Sarah A Sundius, Rachel A Kuske, Sam P. Brown, Edze R Westra

## Abstract

Where there are bacteria, there will be bacteriophages. These viruses are known to be important players in shaping the wider microbial community in which they are embedded, with potential implications for human health. On the other hand, bacteria possess a range of distinct immune mechanisms that provide protection against bacteriophages, including the mutation or complete loss of the phage receptor, and CRISPR-Cas adaptive immunity. Yet little is known about how interactions between phages and these different phage resistance mechanisms affect the wider microbial community in which they are embedded. Here, we conducted a 10-day, fully factorial evolution experiment to examine how phage impact the structure and dynamics of an artificial four-species bacterial community that includes either *Pseudomonas aeruginosa* wild type or an isogenic mutant unable to evolve phage resistance through CRISPR-Cas. Our results show that the microbial community structure is drastically altered by the addition of phage, with *Acinetobacter baumannii* becoming the dominant species and *P. aeruginosa* being driven nearly extinct, whereas *P. aeruginosa* outcompetes the other species in the absence of phage. Moreover, we find that a *P. aeruginosa* strain with the ability to evolve CRISPR-based resistance generally does better when in the presence of *A. baumannii*, but that this benefit is largely lost over time as phage is driven extinct. Combined, our data highlight how phage-targeting a dominant species allows for the competitive release of the strongest competitor whilst also contributing to community diversity maintenance and potentially preventing the reinvasion of the target species, and underline the importance of mapping community composition before therapeutically applying phage.

## Introduction

Microbiome research is a dynamic and growing field in microbiology, producing an incredible amount of sequence data from a wide range of clinical and environmental samples. Humans, for instance, are colonised by a large number of microorganisms and research continues to implicate microbial communities as potential drivers behind multiple important biological processes [1–3]. These processes may play important roles in human health and disease, with some work focusing on correlations based on microbiome composition [4–8] while other look more closely for direct causality [9–12]. Still, the challenge to move beyond descriptive and correlative approaches remains, and there is a need to develop bottom-up mechanistic and quantitative understanding of the forces acting upon and shaping microbial communities. To this end, synthetic polymicrobial communities are being designed, and are gaining traction in both pure and applied microbiome studies [13–16]. Synthetic microbiomes allow for precise and malleable experimental testing of the basic rules that govern both microbial organisation and functioning on molecular and ecological scales [17–20], as well as allowing for exploration of causal roles connecting specific microbiome structures to potential outcomes of interest.

Bacteria and their viral predators, bacteriophages (phages), have long been of interest in microbiological research, in part due to being the most abundant biological entity on the planet [21,22]. Phages are highly diverse in terms of their morphology, genetics, and life histories [21,23], with a clear distinction between obligatory killing lytic phages and temperate phages that can either cause a dormant infection (lysogenic cycle) or cell lysis to release new phage particles (lytic cycle). Phages are thought to play a key role in shaping both the taxonomic and functional composition of microbial communities as well as their stability, ecology and evolution [23–26]. For example, lytic replication will per definition cause a reduction in the density of the bacterial host strain or species, which in turn can have knock-on effects for the microbial community composition through the enabling of invasion and/or co-existence of competitor species. Despite the large potential impact of lytic phage, only a very limited number of experimental studies have explored the ecology and evolution of bacteria-phage interactions in a microbial community context [27,28], and it remains unclear if and how interactions between different species in more complex communities shape the effects of lytic phages on microbial eco-evolutionary dynamics. Consequently, we lack the steppingstones to understand how phages shape microbial community dynamics (reviewed in [23]), which are urgently needed to understand potentially causal relationships between natural phage communities and a variety of human diseases [29–34], and for optimising the therapeutic application of phages in the clinic.

A key consideration in this context is that bacteria can overcome phage infection through a range of different means [35,36], with varied underlying molecular mechanisms and which can act during different stages of phage infection [37–40]. Through the modification, masking or complete loss of phage-binding surface receptors for example, bacteria can prevent phage adsorption and injection [39,41]. Systems such as CRISPR-Cas on the other hand work by inserting short DNA sequences from phage and other invasive mobile genetic elements into the host genome to provide future immunological memory [42]. Unlike CRISPR-based resistance [15], phage resistance through receptor mutation can be associated with substantial phenotypic shifts and fitness trade-offs, through changes to virulence [43,44], biofilm formation [45], or antibiotic resistance [46].

Combining an exploratory and hypothesis driven approach, we applied experimental evolution to examine how a phage impacts the dynamics of an artificial bacterial community. This community consisted of *Pseudomonas aeruginosa*, *Staphylococcus aureus, Acinetobacter baumannii*, and *Burkholderia cenocepacia*, all of which are opportunistic pathogens that can cause severe infection and may co-infect with one another [47–50]. Firstly, we hypothesised that the addition of a *P. aeruginosa* specific phage would promote species coexistence by limiting *P. aeruginosa* dominance through competitive release (expansion of phage resistant competitors, following removal of phage susceptible competitor) in a way akin to what is commonly observed with antibiotics [14,51–54]. Secondly, we hypothesised that blocking the ability of *P. aeruginosa* to evolve CRISPR-based immunity would reduce *P. aeruginosa* persistence due to community dependent fitness costs of surface-modification [15]. We found that the addition of a *P. aeruginosa* targeting phage resulted in the general maintenance of community diversity and coexistence, but also a shift in dominant species from *P. aeruginosa* to *A. baumannii* – with the former being unable to reinvade even after the phage was driven extinct. The impact of the type of phage resistance was limited or transient, however: While a *P. aeruginosa* wild-type with the ability to evolve CRISPR-based phage resistance initially had a slight fitness advantage in the presence of *A. baumannii* over its CRISPR-negative isogenic mutant, this effect was lost over time as the phage was driven extinct.

## Results

To measure the effect of phage on microbial community dynamics, we carried out a fully factorial 10-day *in vitro* evolution experiment using all possible combinations of one, two, three or four competitor species: *S. aureus*, *A. baumannii*, *B. cenocepacia*, and *P. aeruginosa* PA14 in the presence or absence of lytic phage DMS3vir. Additionally, to examine the impact of CRISPR-Cas vs surface modification on these dynamics, we used both the wild-type (WT) *P. aeruginosa* PA14 strain, which can evolve CRISPR-based phage resistance and an isogenic mutant lacking a functional CRISPR system. Following inoculation, we tracked the microbial community dynamics for all experimental treatments at regular intervals over a period of 10 days. All experiments were conducted in Lysogeny Broth (LB) at 37°C (see methods for details).

### *P. aeruginosa* dominates in the absence of phage

Without *P. aeruginosa* present in the community, *S. aureus* was primarily the dominant species – with the ability to co-exist with *A. baumannii* while outcompeting *B. cenocepacia* (Figs 1 and S1). This, however, was not reflected once *P. aeruginosa* was introduced to the community. In the absence of phage, *P. aeruginosa* quickly became the dominant species in the microbial community, regardless of starting composition and the *P. aeruginosa* genotype (PA14 WT vs CRISPR-KO) (Figs 2 and S3). Consistent with this, the densities of the competitor species rapidly declined during these co-culture experiments (Fig 2). Yet there was a clear difference in the rate at which competitor species declined in frequency, which was highest for *S. aureus* and lowest for *A. baumannii* (Fig 2, ANOVA: effect of treatment on *S. aureus*; F = 2.2, p = 0.09; overall model fit; adjusted R^2^ = 0.60, F_20,171_ = 15.45, p < 2.2 x 10^-16^: effect of treatment on *A. baumannii*; F = 0.52, p = 0.67; overall model fit; adjusted R^2^ = 0.66, F_20,171_ = 19.89, p < 2.2 x 10^-16^: effect of treatment on *B. cenocepacia*; F = 1.36, p = 0.26; overall model fit; adjusted R^2^ = 0.69, F_20,171_ = 22.45, p < 2.2 x 10^-16^).

**Fig 1.**
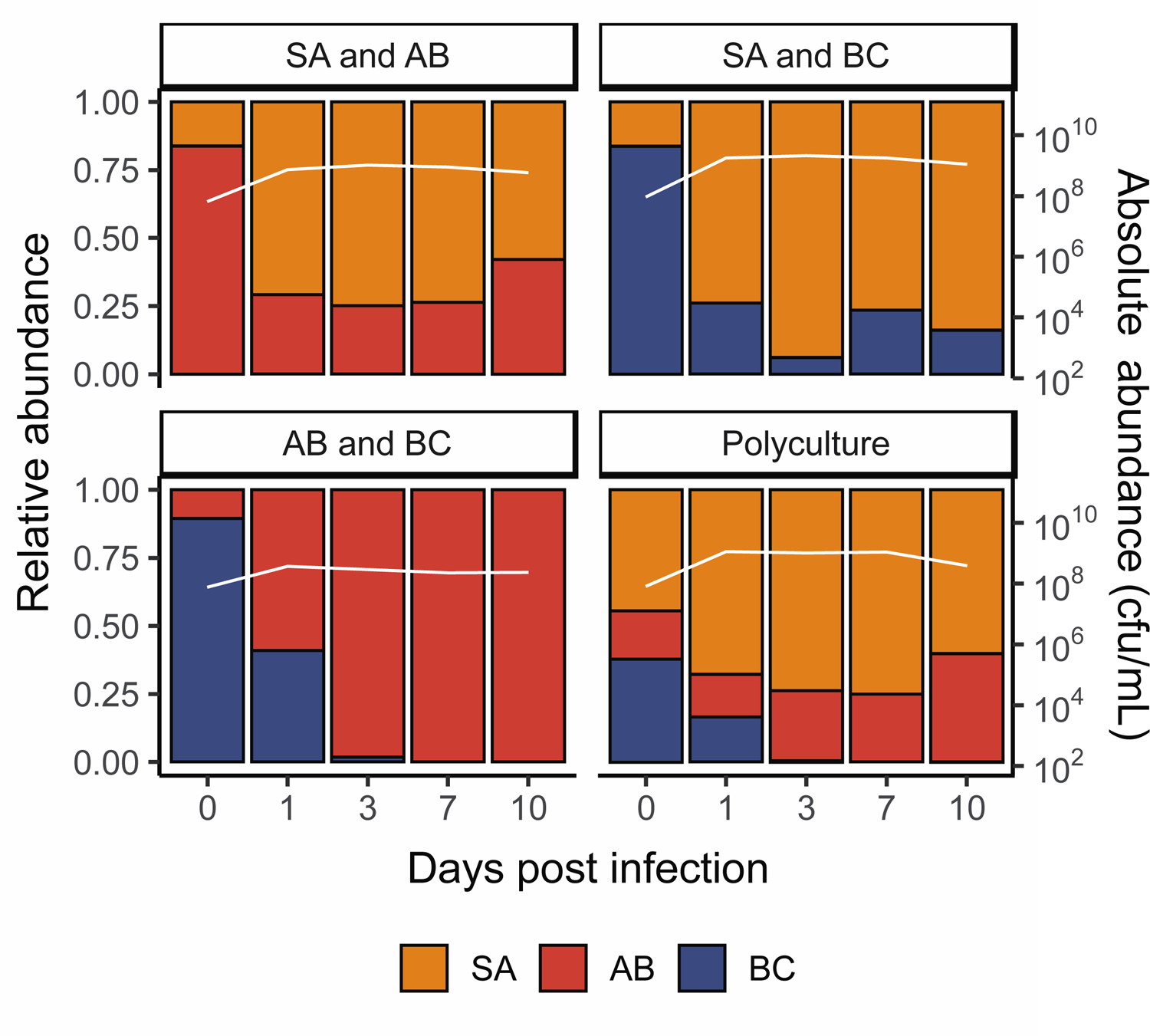
*S. aureus* and *A. baumannii* both perform well in the absence of P. aeruginosa. Showing the community composition and bacterial densities in cfu/mL over time for the microbial communities in the absence of *P. aeruginosa*. The community composition was estimated by qPCR at days 0, 1, 3, 7 and 10 of the experiment. The coloured bars represent the relative abundance of each species (left y axis), while the white line represents total abundance in cfu/mL (right y axis). Each panel represents average composition across six replicates for each treatment over time. SA = *S. aureus*, AB = *A. baumannii*, BC = *B. cenocepacia*. For individual replicates of species abundance, see Fig. S1.

**Fig 2.**
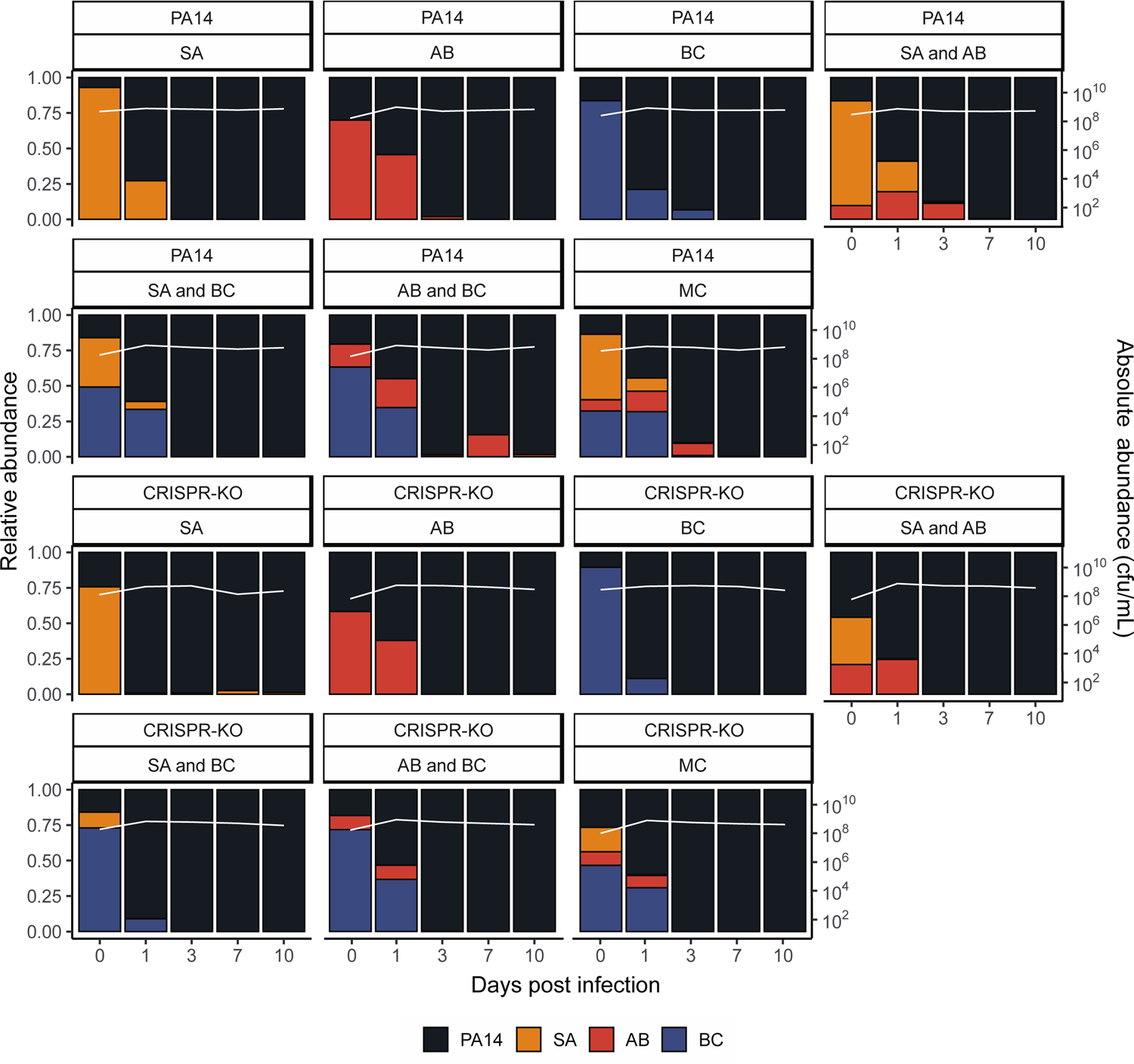
*Pseudomonas aeruginosa* becomes the dominant species in the absence of phage. Showing the community composition and bacterial densities in cfu/mL over time for the microbial communities in the absence of phage. The community composition was estimated by qPCR at days 0, 1, 3, 7 and 10 of the experiment. The coloured bars represent the relative abundance of each species (left y axis), while the white line represents total abundance in cfu/mL (right y axis). Each panel represents average composition across six replicates for each treatment over time. PA14 = *P. aeruginosa*, SA = *S. aureus*, AB = *A. baumannii*, BC = *B. cenocepacia*, MC = microbial community. For individual replicates of species abundance, see Fig. S3.

While the microbial community dynamics were relatively similar for the WT and CRISPR-KO strains, some significant differences were observed. For example, the densities of the CRISPR-KO strain were slightly lower in the presence compared to the absence of *S. aureus* on its own (Fig 2, linear model: t = 2.048, p = 0.0413; overall model fit; adjusted R^2^ = 0.21, F_36,345_ = 3.77, p < 6.03 x 10^-11^). Moreover, *S. aureus* and *A. baumannii* reached higher densities in the presence of the PA14 WT compared to the CRISPR-KO strain, particularly at the earlier timepoints (Fig 2). In contrast to this, densities of *B. cenocepacia* over time were similar in the presence of both *P. aeruginosa* genotypes (Fig 2). Regardless these minor differences, *P. aeruginosa* consistently and readily outcompeted the other community members in the absence of phage, with all three being extinct or close to extinction by day 10 (Fig 2). For visualisation purposes, the data from Figure 2 is also presented as an ordination plot (Fig S2).

### Phage affects microbial community dynamics

Whereas *P. aeruginosa* dominated in the absence of phage, we hypothesised this would change once a PA14 targeting phage (DMS3vir) was introduced, largely by a virulent phage reducing the susceptible host population, facilitating expansion of other species through competitive release [14,51–54]. As expected, phage DMS3vir initially reached high titres due to replication on sensitive *P. aeruginosa* hosts, followed by a rapid decline in phage densities due to the evolution of phage resistance, regardless of whether the host had a functional CRISPR-Cas system or not (Fig S4). Crucially however, the presence of phage caused microbial communities to no longer be dominated by *P. aeruginosa*, as when compared to the no phage treatments, very few to none of the experimental repeats had one or more bacterial species go extinct, with *A. baumannii* reaching particularly high abundance (Figs 3 and S5). It is here worth noting that while *B. cenocepacia* is not visible at later timepoints in the compositional plot due to low relative abundance of <0.1 (Fig 3), we consistently observed persistence of *B. cenocepacia* at an average of ∼10^4^ cfu/mL across all treatments (see Fig S5). For visualisation purposes, the data from Figure 3 is also presented as an ordination plot (Fig S6).

**Fig 3.**
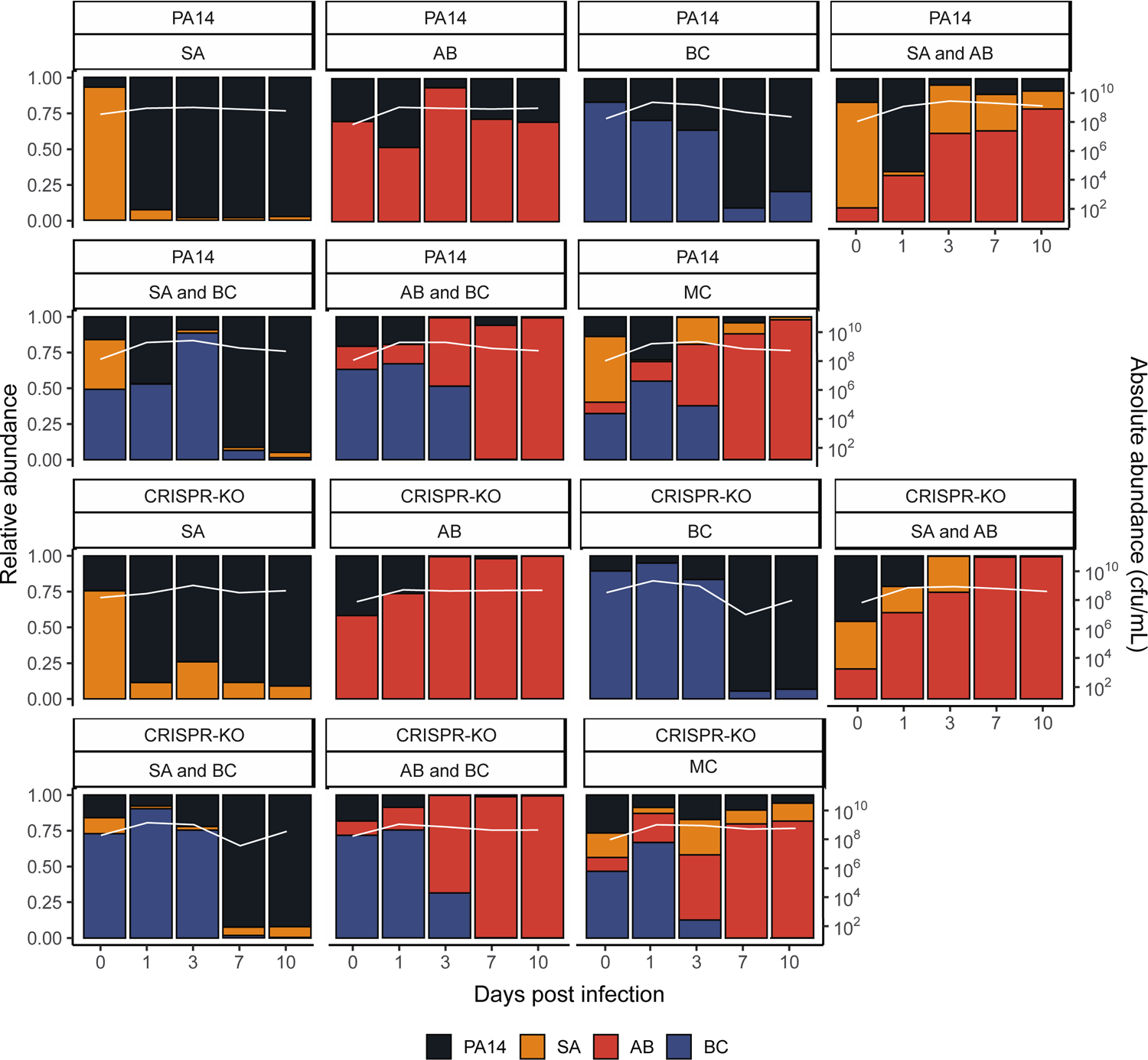
Phage allows for the maintenance of all microbial community members, with *A. baumannii* becoming the new dominant species. Showing the community composition and bacterial densities in cfu/mL over time for the microbial communities in the presence of phage. The community composition was estimated by qPCR at days 0, 1, 3, 7 and 10 of the experiment. The coloured bars represent the relative abundance of each species (left y axis), while the white line represents total abundance in cfu/mL (right y axis). Each panel represents average composition across six replicates for each treatment over time. PA14 = *P. aeruginosa*, SA = *S. aureus*, AB = *A. baumannii*, BC = *B. cenocepacia*, MC = Microbial community. For individual replicates of species abundance, see Fig. S5.

Interestingly, the PA14 WT generally reached greater relative abundance than the CRISPR-KO strain when in the presence of *A. baumannii*, consistently doing so early in the experiment when phage remained in the population (Figs 3 and S2). This was in concordance with *P. aeruginosa* evolving higher levels of CRISPR-based immunity against phage DMS3vir in treatments including *A. baumannii* due to the increased fitness cost of surface modification (Fig S7 and [15]): At 3 days post infection, there was a significant effect of all treatments on the proportion of CRISPR-based resistance that had evolved compared to the PA14 monoculture, but this effect was strongest for treatments that contained *A. baumannii*. At timepoint 10 we only found an increased proportion of *P. aeruginosa* clones immune through CRISPR-Cas when the treatment included *A. baumannii* (GLM; *A. baumannii*; t = 2.637, p = 0.01; *S. aureus* and *A. baumannii*, t = 2.283, p = 0.025; A. *baumannii* and *B. cenocepacia*, t = 2.689, p = 0.0087; polyculture, t = 2.141, p = 0.035).

### The type of evolved phage resistance does not have a knock-on effect on microbial community dynamics

We have previously sown that the evolution of phage resistance by mutation of the Type IV pilus (the phage receptor) is associated with large fitness trade-offs in a microbial community context, whereas evolution of CRISPR-based immunity is not associated with any detectable trade-offs [15]. We therefore predicted that the ability to evolve phage resistance through CRISPR-Cas would also have knock-on effects for the microbial community dynamics. However, measurement of the abundance of the competitors revealed that these were overall largely unaffected by the presence of a functional CRISPR-Cas immune system in *P. aeruginosa* with the exception of *S. aureus*: In the presence of the *P. aeruginosa* WT strain, *S. aureus* densities were significantly lower in two of the microbial communities compared to the same co-culture experiments with the CRISPR-KO strain (Figs 3 and S5, Effect of *P. aeruginosa* clone on *S. aureus* abundance, linear model: Treatment *S. aureus*; t = −2.363, p = 0.0216, adjusted R^2^ = 0.2659, F_14,57_= 2.837, p = 0.002786; Treatment *S. aureus* and *A. baumannii*; t = −2.043, p = 0.0457, adjusted R^2^ = 0.3867, F_14,57_= 4.198, p = 5.3 x 10^-5^).

### A *P. aeruginosa* targeting phage results in the competitive release of *A. baumannii* and general diversity maintenance

We hypothesised that the effect of phage on microbial community structure could largely be explained by the competitive release (increase in absolute abundance, following removal of competitor) of *A. baumannii*, which then takes over to become the dominant species [55]. To assess this, we examined the fold change difference for the final abundance of all three community members in the presence versus absence of phage (Fig 4). Crucially, this revealed a strong increase in *A. baumannii* density in the presence of a phage, supporting the idea that it becomes the dominant and determinant community member when *P. aeruginosa* is inhibited by phage (Fig 3). By contrast, when phage was added, *S. aureus* only experienced a clear fold change increase if it was co-cultured with the CRISPR-KO strain and an additional competitor species. *B. cenocepacia* meanwhile seemed to be the species with the least benefit of phage, but still with a small fold change increase for some treatments (Fig 4).

**Fig 4.**
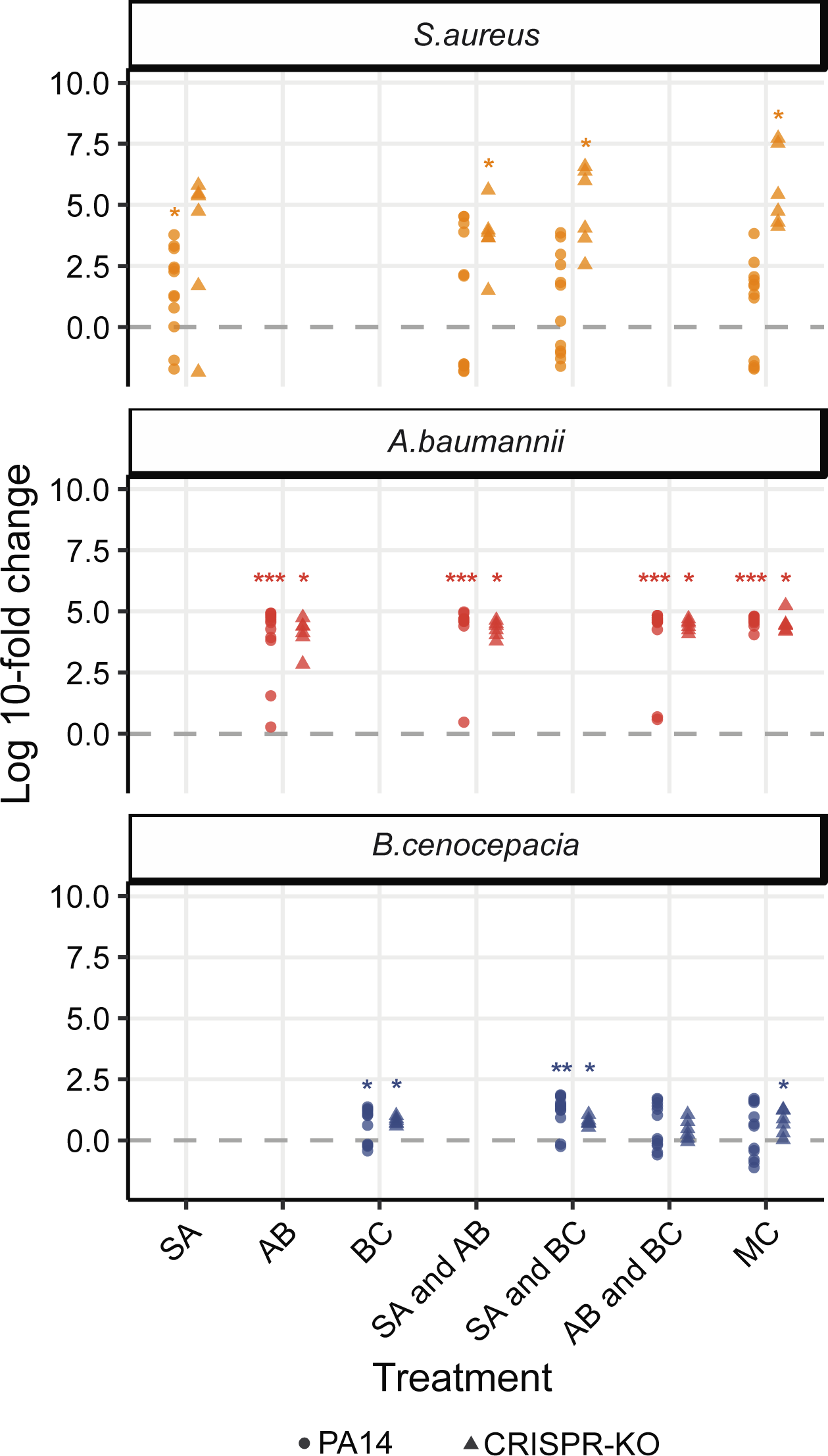
Fold change between no phage and phage treatments at the end of the experiment. The fold change difference of the individual community species not targeted by phage when comparing absolute densities in the presence of phage to the absence at the final experimental timepoint. Asterisks indicate higher final absolute density in the presence versus absence of phage (Wilcoxon signed rank exact test: * p < 0.05, ** p < 0.01, *** p < 0.001).

The substantial 5-fold increase in *A. baumannii* given the presence of phage (Fig 4) reflects a sustained divergence in the trajectory of *A. baumannii* in the phage treatments, despite the attenuation of phage titre by day 7 (Fig S4). We hypothesised that the lack of *P. aeruginosa* rebound after phage clearance was due to a frequency-dependent shift in competitive dominance. To test this hypothesis, we competed ancestral *A. baumannii*, *S. aureus* and *B. cenocepacia* against increasingly rare *P. aeruginosa* challenge, and found no barrier to *P. aeruginosa* invasion in pairwise experiments, down to a frequency of 1 in 10,000 cells (Fig 5). This result suggests that the failure of *P. aeruginosa* to return to dominance following phage clearance is due to more complex community-mediated interactions.

**Fig 5.**
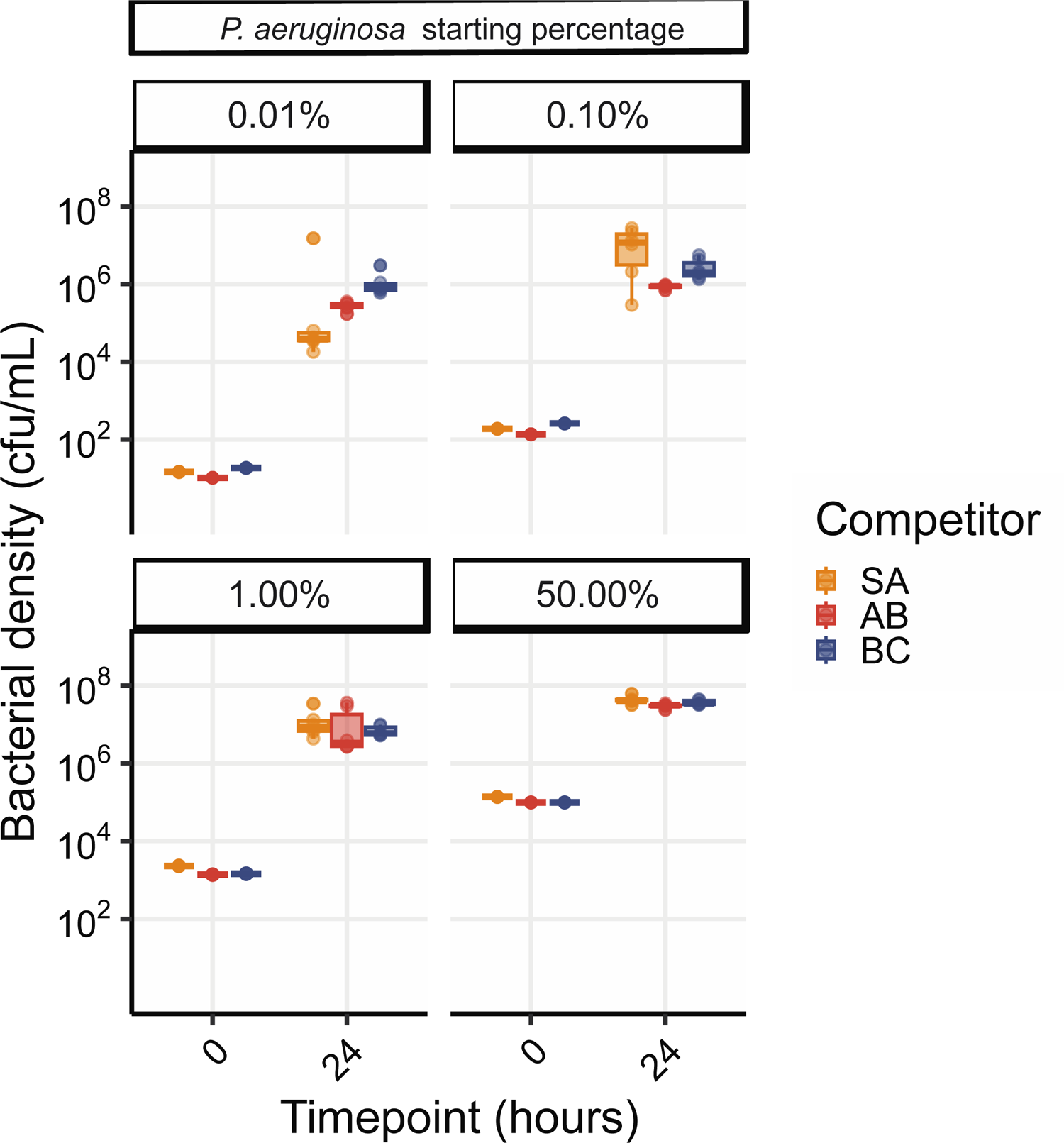
*P. aeruginosa* can reinvade from rare against all community members. Showing *P. aeruginosa* density in cfu/mL from competition experiments between PA14 wild-type with variable starting densities against either *S. aureus* (SA), *A. baumannii* (AB) or *B. cenocepacia* (BC). The species densities were estimated by qPCR at time-point 0 and 24 h post co-culture. Box plots show the median, 25th and 75th percentile, and the interquartile range. Raw values from each replicate are shown as points (n= 6 per pairwise competition).

To quantitatively assess changes in community diversity, we calculated Shannon diversity indexes for all experimental treatments. We hypothesised that the addition of phage not only results in competitive release of one other bacterium (Fig 4), but facilitates general maintenance of microbial diversity. Plotting these diversity scores over time shows that without phage there is a rapid loss of diversity over time, whereas community complexity persists in the presence of phage (Fig 6: ANOVA: PA14 WT effect of phage; F = 27.57, p = 2.3 x 10^-7^; CRISPR-KO effect of phage; F = 89.19, p < 2.2 x 10^-16^; Overall model fit for PA14 WT: adjusted R^2^ = 0.64, F_38,465_ = 24.87, p < 2.2 x 10^-16^; Overall model fit for CRISPR-KO: adjusted R^2^ = 0.56, F_32,303_ = 14.56, p < 2.2 x 10^-16^). This was true for treatments for both *P. aeruginosa* genotypes, but the trend became most pronounced for the CRISPR-KO strain when applying direct comparisons using Tukey contrasts, in which case we found phage to significantly increase diversity over time in nearly all treatments (Fig 6, indicated by asterisks).

**Fig 6.**
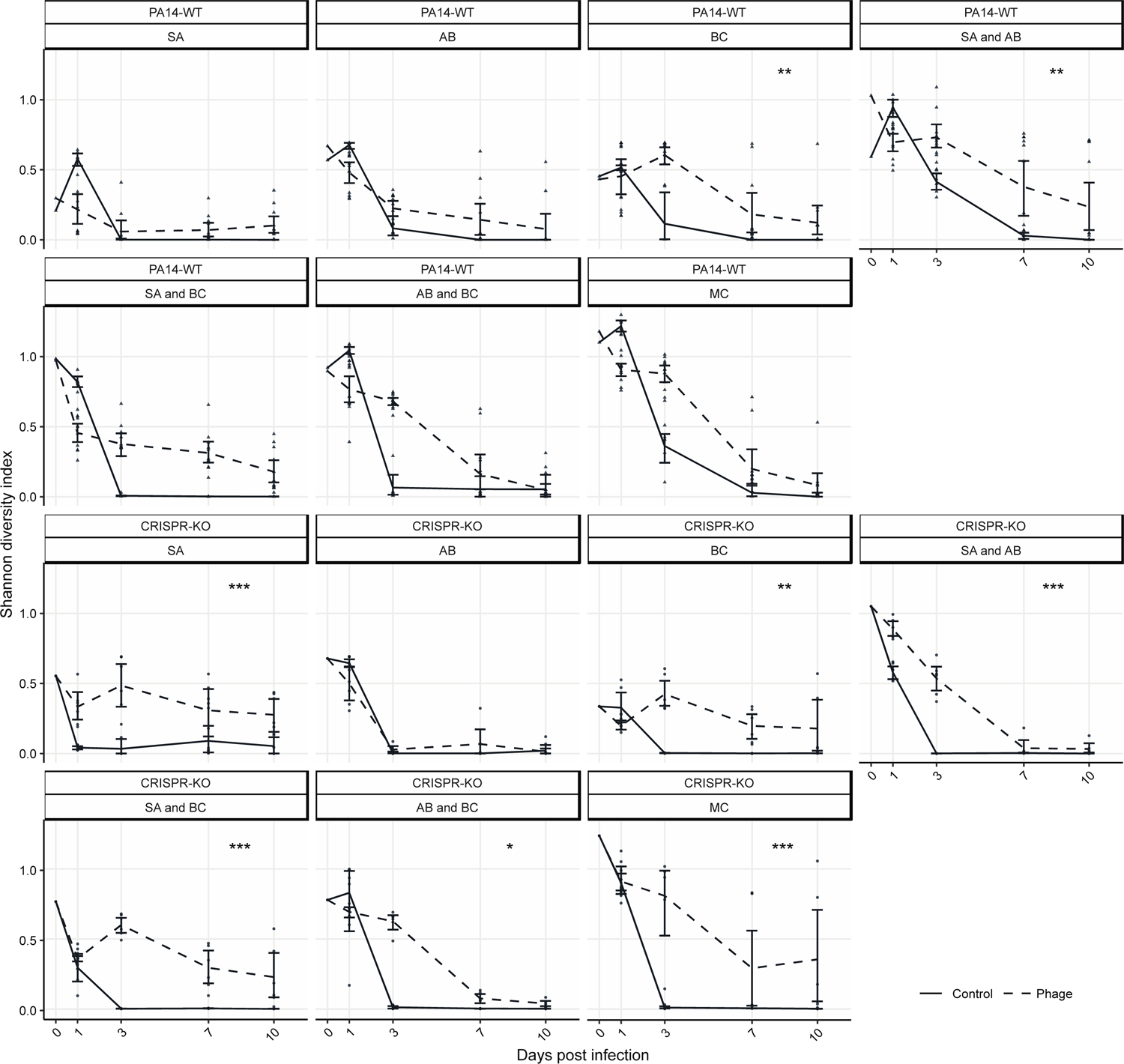
Shannon diversity over time illustrating the diversity maintaining effects of phage. The change in diversity over time, illustrated using Shannon diversity indexes, for both the PA14 WT and CRISPR-KO strains across all treatments (SA = *S. aureus*, AB = *A. baumannii*, BC = *B. cenocepacia,* MC = microbial community). Data are mean ± 95% CI, and asterisks indicate a significant difference over time in Shannon diversity between treatments with phage or no phage (n = 6 per timepoint for all expect the PA14 WT with phage treatments, where n = 12) (effect of *P. aeruginosa* clone; linear model with Tukey contrasts: * p < 0.05, ** p < 0.01, *** p < 0.001).

### Four species community dynamics are predictable from two and three species community data, in the absence of phage

A major challenge in synthetic community research is developing robust modelling frameworks that are capable of predicting community dynamics [56]. In a final set of analyses, we sought to assess the predictive performance of generalized Lotka Volterra (gLV) competition equations, trained on just 2-species data or a combination of 2- and 3-species data. Our results showed that fitting gLV models with pairwise only datasets led to predictive failures when applied to 3- or 4-species datasets (Fig S8), consistent with the presence of higher order interactions effects (when the effect of species A on species B is dependent on the presence of species C [57]). In contrast, fitting gLV models to 2- and 3-species data and using the resulting interaction terms to predict 4-species dynamics reasonably fit the data in the absence of phage (Fig 7).

**Fig 7.**
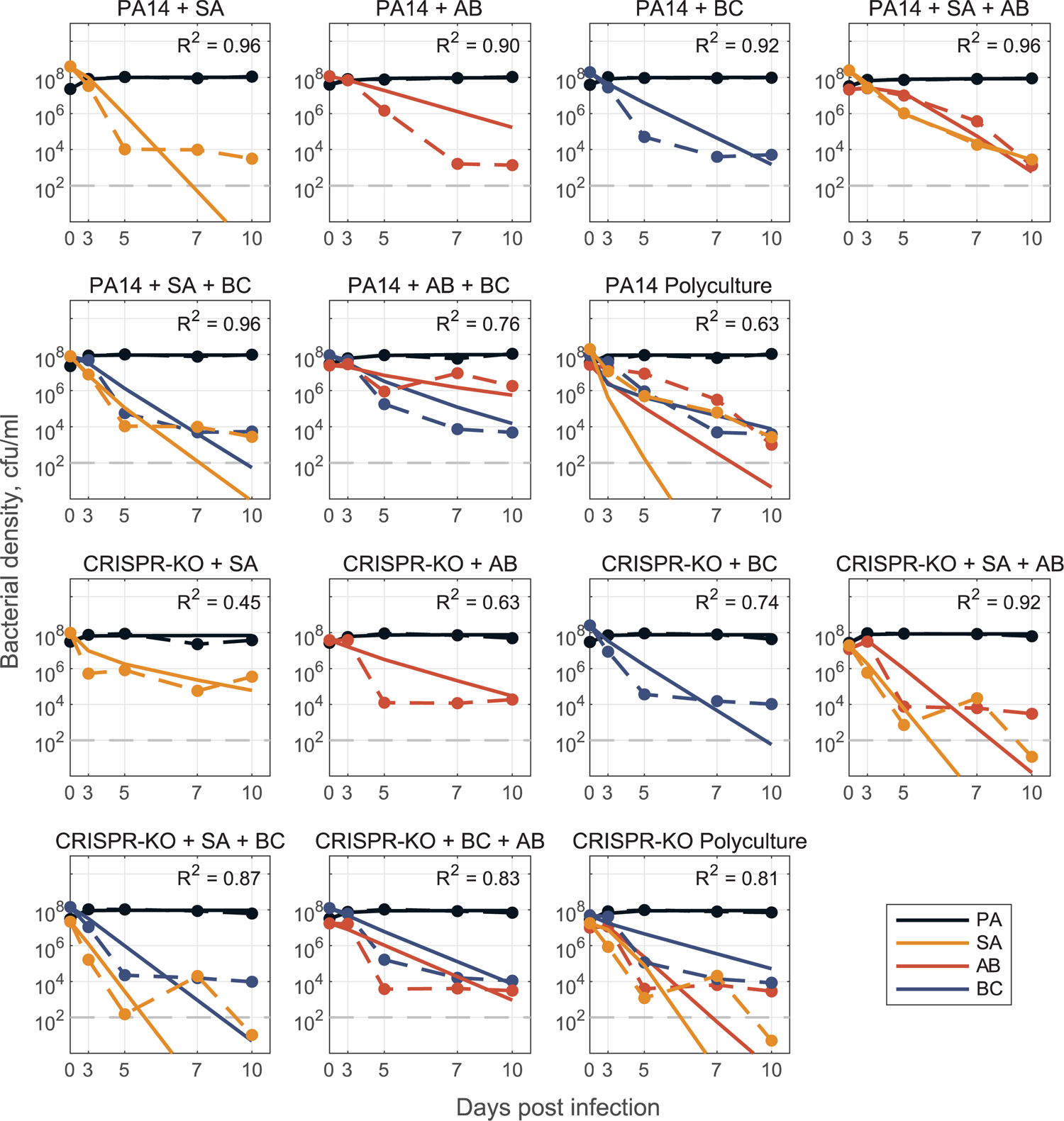
Model for no phage data. Model fit predictions for two-, three-, and full four species community dynamics (solid lines) compared to experimental data (dashed lines). Models of 2- and 3-species dynamics were parameterized via optimization with least-squares to fit to a system of ODEs (defined as a generalized Lotka-Volterra competition model with n species, where n=1,2,3,4). Only single species maximal growth rates (*r*_*i*_ for species *i* = 1,…, *n*) were fixed from fitting mono-culture data, all interaction coefficients (*β*_*i*,*j*_ describing the inhibitory effect of species *j* on species *i* for all *i*,*j* = 1,2 in the 2-species case, for all *i*,*j* = 1,2,3 in the 3-species case) were open for fitting. We construct the full 4-species community interaction matrices (one for PA14 and one for CRISPR-KO) by averaging corresponding *β*_*i*,*j*_ interaction terms from the fit 2- and 3-species models (see Text S1), and use this matrix to simulate dynamics in the respective polyculture cases. See Methods and Text S1 for detailed description of mathematical modelling.

In the presence of phage (Fig 8), we again utilised the gLV framework where the impact of phage is implicit (quantified by how interaction coefficients change as compared to the no-phage case). The gLV model framework could adequately describe 2- and 3-species data, but the interaction coefficients did not generalise quantitatively to 4-species data – likely reflecting the structural limitation of a gLV competition model that does not explicitly capture phage predation dynamics. However, the model parameterised with 1-, 2- and 3-species data did capture a qualitative shift in ecological outcomes from sole *P. aeruginosa* survival to competitive release of *A. baumannii* and *S. aureus* when *P. aeruginosa* is targeted by phage (Fig S9).

**Fig 8.**
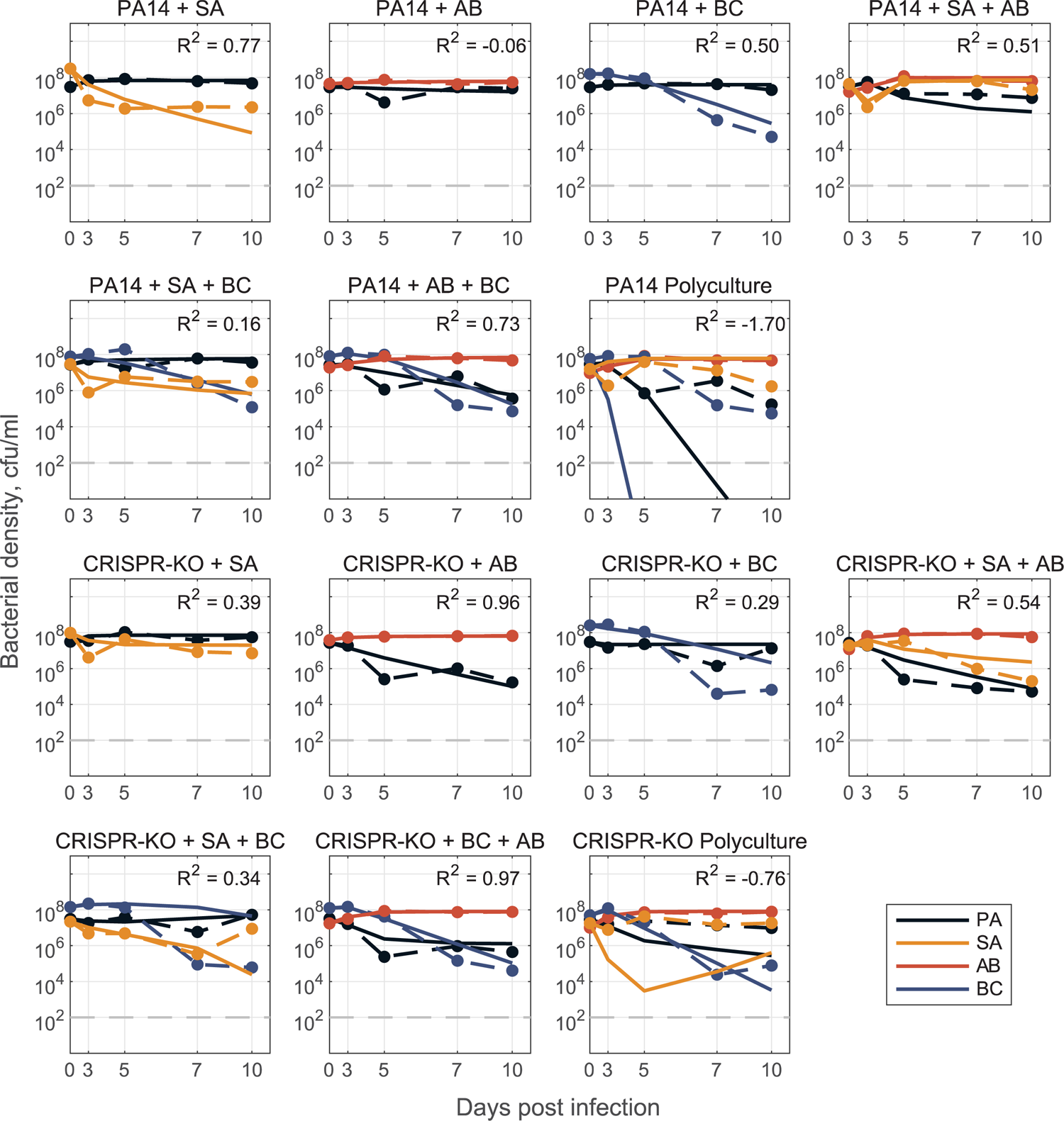
Model for phage data. Model fit predictions for two-, three-, and full four species community dynamics (solid lines) in the presence of phage compared to experimental data (dashed lines). Here, models were parameterized via optimization with least-squares to fit a system of ODEs (defined as a generalized Lotka-Volterra competition model with n species, where n=1,2,3,4), where we don’t explicitly track the phage population dynamics. Instead, we assume that the phage acts as some external perturbation that leads to changes in the interactions between community members (*β*_*i*,*j*_ values differ from values in Figure 7). Only single species maximal growth rates were fixed from fitting mono-culture data (*r*_*i*_ for species *i* = 1,…, *n*), all interaction coefficients were open for parameterizing 2- and 3-species models (*β*_*i*,*j*_ describing the inhibitory effect of species *j* on species *i* for all *i*,*j* = 1,2 for the 2-species case, for all *i*,*j* = 1,2,3 for the 3-species case). We then construct the full 4-species community interaction matrices (one for PA14 and one for CRISPR-KO) by averaging corresponding *β*_*i*,*j*_ interaction terms from the fit 3-species models (treatments: PA+AB+SA, PA+BC+SA, PA+AB+BC with phage), and use this matrix to simulate dynamics in the respective polyculture cases. See Methods and Text S1 for detailed description of mathematical modelling.

## Discussion

The advent of deep sequencing has dramatically increased our knowledge of the composition and functioning of microbiomes both in and around us. The role of microbial communities in human health has consequentially received increasing attention, with research focusing on how changes in microbiome composition over time may affect human health and define patient outcomes (reviewed in 57). In addition, an increasing number of correlational studies find associations between virome composition and the health status of their host [23,59–61], likely mediated by changes in the microbiome that could be either cause or effect. A deeper understanding of the impact of phages on microbiomes is likely to help to infer causal relationships between viromes and human health, and to design optimal therapeutic phage interventions (phage therapy).

Here, we expanded on our previous work on how interspecific competition can shape the evolution of phage resistance in a focal species (*P. aeruginosa*)[15], to study how the interaction between phage and bacterial immune mechanisms affects the broader microbial community dynamics. We found that whereas *P. aeruginosa* dominated in the absence of a phage, the presence of phage resulted in microbial diversity maintenance and *A. baumannii* becoming the dominant species (Figs 3 and 4). Interestingly, the competitive release of *A. baumannii* occurred in all treatments and was virtually independent of whether *P. aeruginosa* had a functional CRISPR-Cas immune system or not. This showed that the amplification of the fitness cost of *P. aeruginosa* receptor mutation in the presence of competitor species [15] has limited impact on the overall community dynamics. Overall, our experimental data align with the notion of phages having the potential to increase microbiome stability [62,63], and support the idea that phages can be useful in the designing of synthetic microbial communities [64]. Surprisingly, our data do not support the hypothesis that bacterial adaptive immune systems play an important role in phage-mediated microbial community structuring under the experimental conditions tested here.

Our mathematical analyses focused on the ability of generalised Lotka-Volterra (gLV) models to predict community dynamics. While our analyses showed reasonable predictive success when incorporating 3-species data, we note that our analyses pose two distinct questions: (1) how can we provide more accurate predictions? (2) what general lessons can we draw from our model analyses?

In agreement with a growing number of gLV-based analyses, we found that a simple ‘bottom up’ model fitting approach (fitting single species growth, then all pairwise interactions, then predicting larger system behaviour [14]) performed poorly, indicating the presence of significant higher order interactions [57,65,66]. Consistent with this conclusion, we found that allowing pairwise interactions to vary (contingent on the presence of a third species) produced both qualitative and quantitative improvements in predicting community dynamics (Figs 7 and 8). In the presence of phage, our model successfully predicted the qualitative result of *A. baumannii* competitive release, but failed to quantitatively replicate observed community dynamics (Fig 8). This quantitative failure suggests that our underlying gLV model structure excludes critical components, such as higher order and/or heterogeneous (in time or space) interactions as well as the explicit predatory effect of phage on *P. aeruginosa* (also likely time and spatially dependent). Additionally, it emphasises an ongoing need in microbiome modelling to evaluate functional forms that can efficiently – with respect to parameter number – and accurately capture the complexities of community dynamics.

Our parameterised models are tuned to the data generated by our specific 4-species community, which raises the question of ‘can we learn more general lessons from our model?’ If we simplify our analysis to a 2-species context (focal pathogen, subject to phage, plus a second, non-focal species), we can translate recent analyses on the impact of (antibiotic) perturbations in a two species context [54]. This approach delivers a couple of general messages. First, we can provide a general mathematical definition of ‘competitive release’ mediated by phage predation (see Text S1) highlighting the importance of both demographic and species interaction parameters. Second, we can underline that phage control of a focal pathogen presents secondary ecological problems, if the pathogen is competing with other pathogens that are not targeted by the phage. In this scenario, phage therapy (or other ‘narrow spectrum’ treatment) can lead to competitive release of previously rare pathogens, as seen in our experimental data showing the replacement of *P. aeruginosa* by *A. baumannii*, following phage treatment. These results imply that ‘narrow spectrum’ anti-microbials, such as phages, may not always be the best option when multiple pathogen species are competing within a single polymicrobial infection. One counter-intuitive suggestion, grounded in the idea of ‘beneficial resistance’ [54], is to co-administer probiotic competitors that are resistant to the treatment (i.e. phage or antibiotic resistant) and can therefore continue to exert ecological suppression on the focal pathogen during the course of treatment, while presenting minimal direct risk of disease. Alternatively, one could apply phage cocktails that target not just the dominant pathogen, but also other co-existing bacterial pathogens, to pre-emptively prevent their invasion [67].

## Materials and Methods

### Bacteria and phages

The bacteria *P. aeruginosa* UCBPP-PA14 strain marked with streptomycin resistance, the PA14 *csy3::LacZ* strain (CRISPR-KO), and phages DMS3vir and DMS3vir+acrF1 were used throughout this study and have all been previously described [67,68]. The microbial community consisted of *S. aureus* strain 13 S44 S9, *A. baumannii* clinical isolate

FZ21, and *B. cenocepacia* J2315, and were all isolated at Queen Astrid Military Hospital, Brussels, Belgium.

### Evolution experiment

The evolution experiment was performed by inoculating 60 µl from overnight cultures, that were grown for 24 hours, into glass microcosms containing 6 ml fresh LB medium (60 µl of culture containing ca. 10^6^ cfu). All polyculture mixes were prepared so that *P. aeruginosa* made up approximately 25% of the total inoculation volume (15 µl of 60 µl), with the rest being made up of one or equal amounts of the microbial community bacteria. In all monoculture controls, *P. aeruginosa* was diluted in LB medium to adjust starting densities for consistency across all treatments (*n* = 6 per treatment, unless indicated otherwise). Phage DMS3vir was added at 10^6^ PFU. prior to inoculation. The experiment ran for ten days, with transfers of 1:100 into fresh LB medium being done every 24 hours. Throughout the experiment, the bacterial mixtures were grown at 37°C and shaking at 180 r.p.m. Phage titres were monitored daily, and were determined using chloroform-treated lysate dilutions which were spotted onto lawns of *P. aeruginosa csy::LacZ*. To determine which mechanism of phage-resistance had evolved, 24 randomly selected clones per treatment replica from timepoints 3 and 10 were analysed using methods as detailed in Westra *et al*. 2015 [68].

### DNA extraction and qPCR

Bacterial densities, for both PA14 strains and the other individual microbial community bacteria, were determined using DNA extractions followed by qPCR analyses. DNA extractions were done using the DNeasy UltraClean Microbial Kit (Qiagen), following instructions from the manufacturer, but with an additional pre-extraction step where samples were treated with 15 µl lysostaphin (Sigma) at 0.1 mg ml^-1^ as previously described [15] to ensure lysis of *S. aureus*. The qPCR primers for *P. aeruginosa*, *A. baumannii*, and *B. cenocepacia* were the same as in Alseth *et al*. [15], whereas the *S. aureus* primers used are previously described [69]. All reactions were done in triplicates, using Brilliant SYBR Green reagents (Agilent) and the Applied Biosystems QuantStudio 7 Flex Real-Time PCR system. For reaction mixture and details on PCR programme, see ref. [15]. Bacterial cfu/ mL were calculated from the quantities offered by the standard curve method, adjusting for gene copy number (4, 1, 6, and 6, for *P. aeruginosa*, *S. aureus*, *A. baumannii*, and *B. cenocepacia* respectively).

### Competition experiment

All strains were grown overnight at 37°C with agitation in 30 ml glass universals containing 6 ml of LB medium. For pairwise competition assays, bacteria from overnight cultures were mixed thoroughly at different starting densities of PA14 (i.e., for 50% starting density o *P. aeruginosa* - 30 µl of PA14 + 30 µl of competitor strain) and a total of 60 µl inoculated into 6 ml of LB (*n* = 6 per treatment). Bacteria were grown for 24 hours in a shaking incubator at 180 r.p.m at 37°C. Samples of 500 µl were taken at 0 and 24 hours post competition and mixed with equal volume of 60% glycerol and stored at −70°C until further DNA extraction and qPCR analysis to quantify species densities.

To determine the competitive performance of the focal species relative to competitor strain we used the selection rate (r) formula as it follows: (r) each day (t) 1 (n) day post incubation (r = (ln [density strain A at t_n_/density strain A at t_n−1_] – ln [density strain B at t_n_/density strain B at t_n−1_])/day) [70,71].

### Mathematical modelling

Models were parameterized via optimization with least-squares regression to fit the generalized Lotka-Volterra competition model, 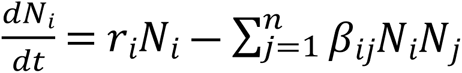, where *N*_*i*_(*t*) is the density of the *i*th species, *r*_*i*_ is the respective single species maximal growth rate, *β*_*ij*_ describes the per capita effect of species *j* on species *i*, and *n* is the total number of species. We take a ‘bottom up’ approach [72] to determine the interaction coefficients *β*_*i*,*j*_. In all cases, we determine single species maximal growth rates *r*_*i*_ from mono-culture time series data and fix them for 2-, 3-, and 4-species model parameterization. Initially, we fit pairwise interaction coefficients for all possible 2-species co-cultures and from here, construct an interaction matrix to predict the dynamics for the 3- and 4-species communities (Fig S8). This is done for both PA14 and CRISPR-KO strains, with and without phage, where phage effects are implicitly represented by changes in interaction parameters between the models with and without phage. To improve results, we additionally fit pairwise interaction parameters *β*_*i*,*j*_ using 3-species experimental data where all interaction parameters are open (only growth rates fixed). Using either the resulting interaction terms or averaging these coefficients with the 2-species coefficients (in PA14 no phage case, Text S1), we are again able to construct an interaction matrix to predict 4-species community dynamics (Fig 7 and 8).

See Text S1 for further description of above model parameterization methods, simulation methods (Fig S9), and mathematical analysis of phage dependent competitive release. All modelling and analysis was done using Matlab 2021b and the code is publicly available at: https://github.com/GaTechBrownLab/phage-community-dynamics.git.

### Statistical analyses

Analysis of the effects of the various species compositions on *P. aeruginosa* densities in the absence (Fig 2) or presence (Fig 3) of phage were done using a generalised linear model (GLM) approach, with log10 cfu/mL set as the response variable. The explanatory variables used in the analyses were type of PA14 clone (PA14 WT or CRISPR-KO), treatment, timepoint, replica, and experimental repeat to account for potential pseudo-replication.

To explore the impact of interspecific competition on the evolution of phage resistance at timepoints 3 and 10 (Fig S7), we used a quasibinomial GLM where the proportion of evolved CRISPR-based phage resistance was the response variable, and treatment, replica, and experimental repeat were the explanatory variables.

The analyses of fold-changes to assess competitive release by comparing absolute density differences of the individual community members in the absence v presence of phage (Fig 4; *S. aureus*, *A. baumannii*, and *B. cenocepacia*) was done through Wilcox signed rank exact tests. A non-parametric test was chosen after performing a Shapiro-Wilk test for normality.

Next, the diversity maintaining effects were examined through assessing the effect of phage DMS3vir on Shannon Diversity index scores over time (Fig 5). This was done through a linear model where the Shannon Diversity index score (H) was the response variable, and treatment, timepoint, the presence of phage, PA14 clone (PA14WT and CRISPR-KO), experimental repeat, and replica were the explanatory variables. Shannon Diversity (H), was calculated as H = -Σ*p*_i_ * ln(*p*_i_), where Σ is the sum and *p*_i_ is the proportion of the entire community made up of species *i*.

For the competition assay (Fig 5), Graphpad Prism9 software (San Diego, CA) was used for statistical analysis. We used one-way ANOVA with Tukey post hoc testing for multiple comparisons, in which, p < 0.05 was considered statistically significant.

Throughout the paper, pairwise comparisons were done using the Emmeans package [73], and model fits were assessed using Chi-squared tests and by comparing Akaike information criterion (AIC) values, as well as plotting residuals and probability distributions using histograms and quantile-quantile plots (Q-Q plots) respectively. All statistical analyses were done using R version 4.3.0. [74], its built-in methods, and the Tidyverse package version 2.0.0 [75]. All data is available at: 10.6084/m9.figshare.24187284.

## Acknowledgements

This work was supported by a grant from the ERC (ERC-STG-2016-714478 - EVOIMMECH), a Biotechnology and Biological Sciences Research Council (BBSRC) sLoLa grant BB/X003051/1, awarded to E.R.W, a NHI grant (NIH 5R21AI156817-02) awarded to S.P.B, and a BBSRC-National Science Foundation (NSF) grant xxxx, awarded to S.P.B., R.A.K & E.R.W.

---

**Supplemental Fig 1.**
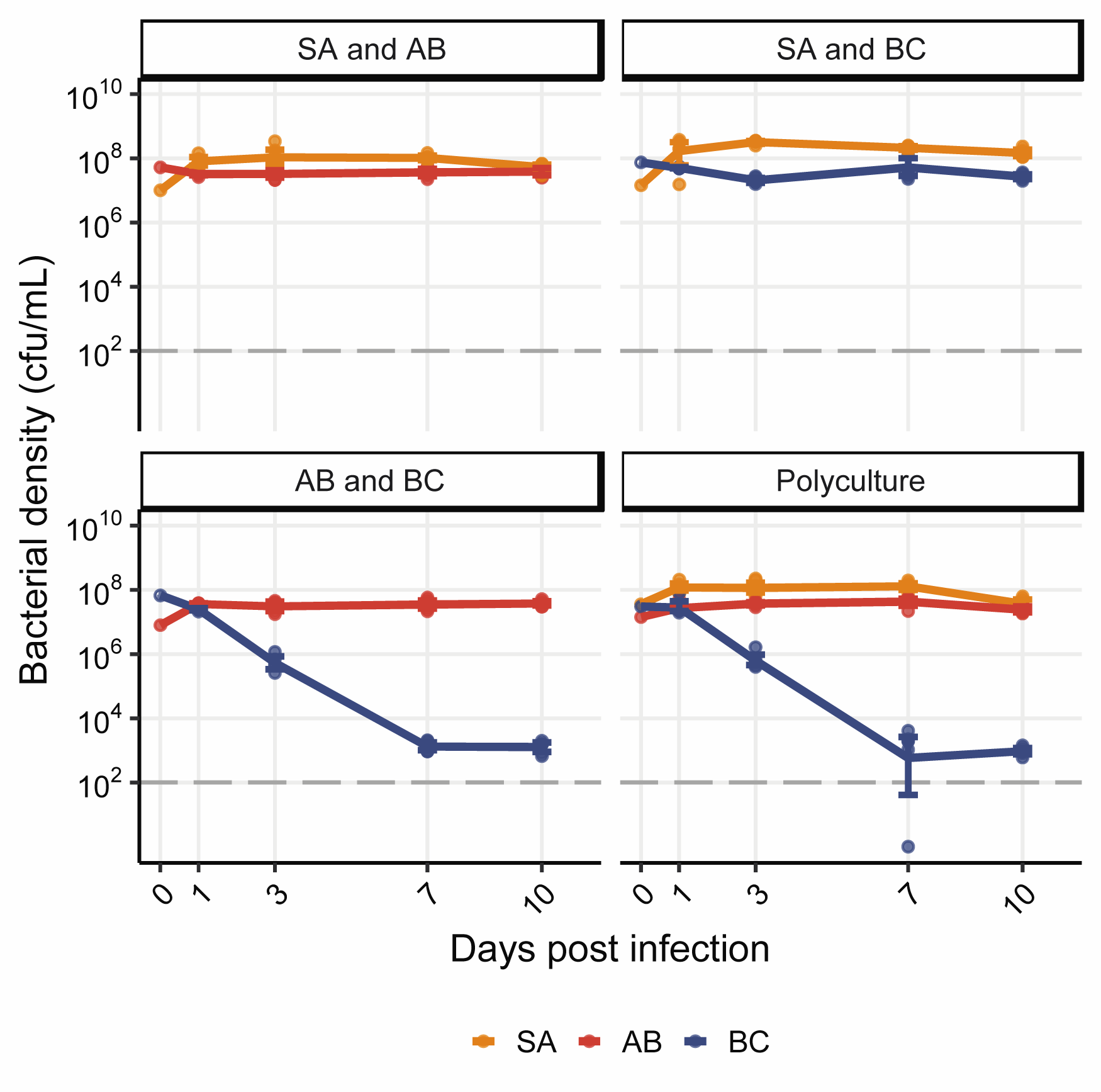
Line plot of bacterial densities in the absence of *P. aeruginosa* and its phage. Showing the bacterial densities in cfu/mL over time for SA (*S. aureus*), AB (*A. baumannii*), and BC (*B. cenocepacia*) in various co-culture combinations in the absence of *P. aeruginosa* and its phage. Dashed horizontal line at 10^2^ cfu/mL marks the threshold of reliable detection where the qPCR results indicate the bacteria has gone or is close to extinction from a population. Data are mean ± 95% CI.

**Supplemental Fig 2.**
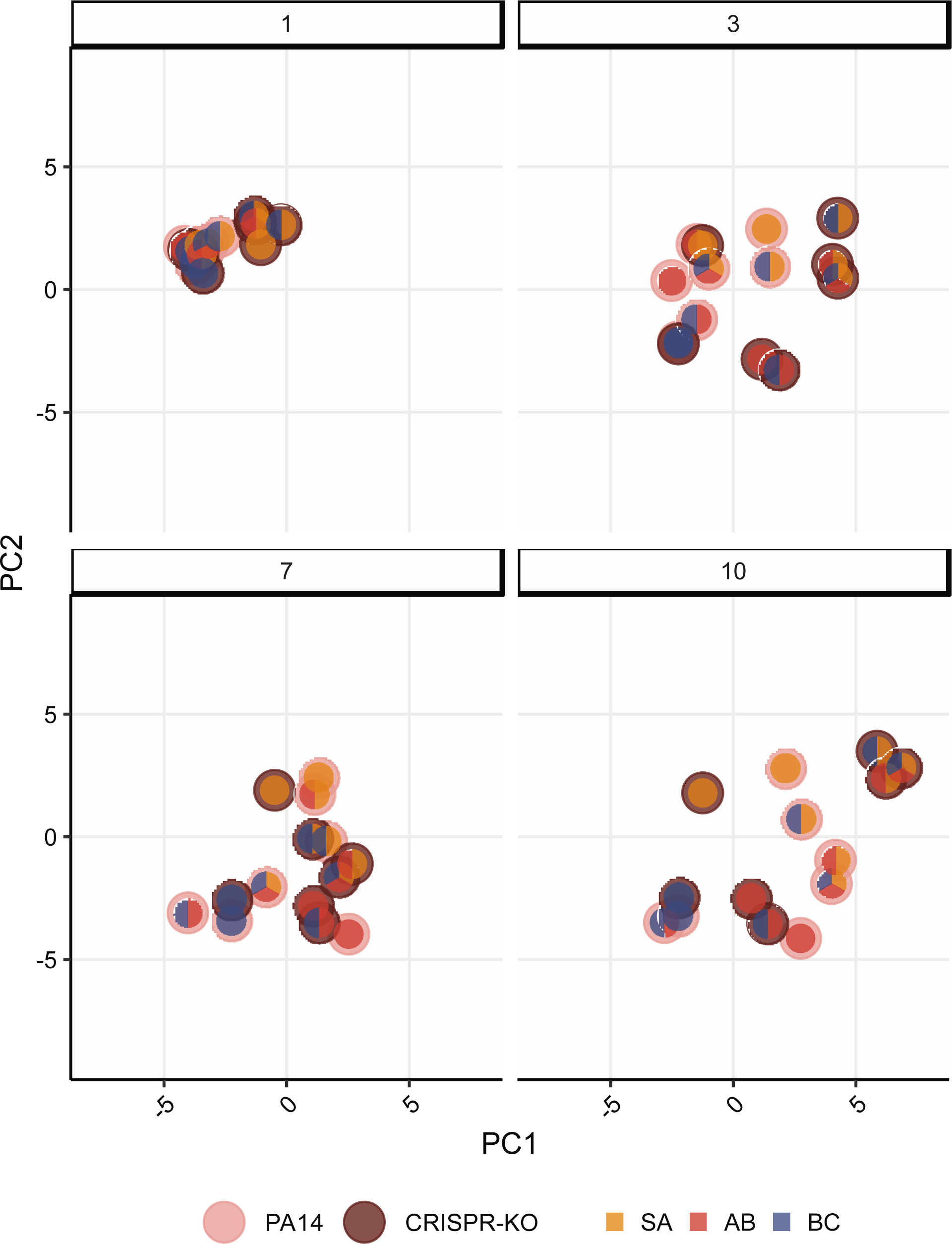
Ordination plot in the absence of phage. PCA ordination of relative bacterial abundance in the absence of phage DMS3vir, with grid layouts separated into days post phage infection. Outer circle colour indicates which PA14 clone is present in the population, while inner circle indicates community composition (SA = *S. aureus*, AB = *A. baumannii*, BC = *B. cenocepacia*).

**Supplemental Fig 3.**
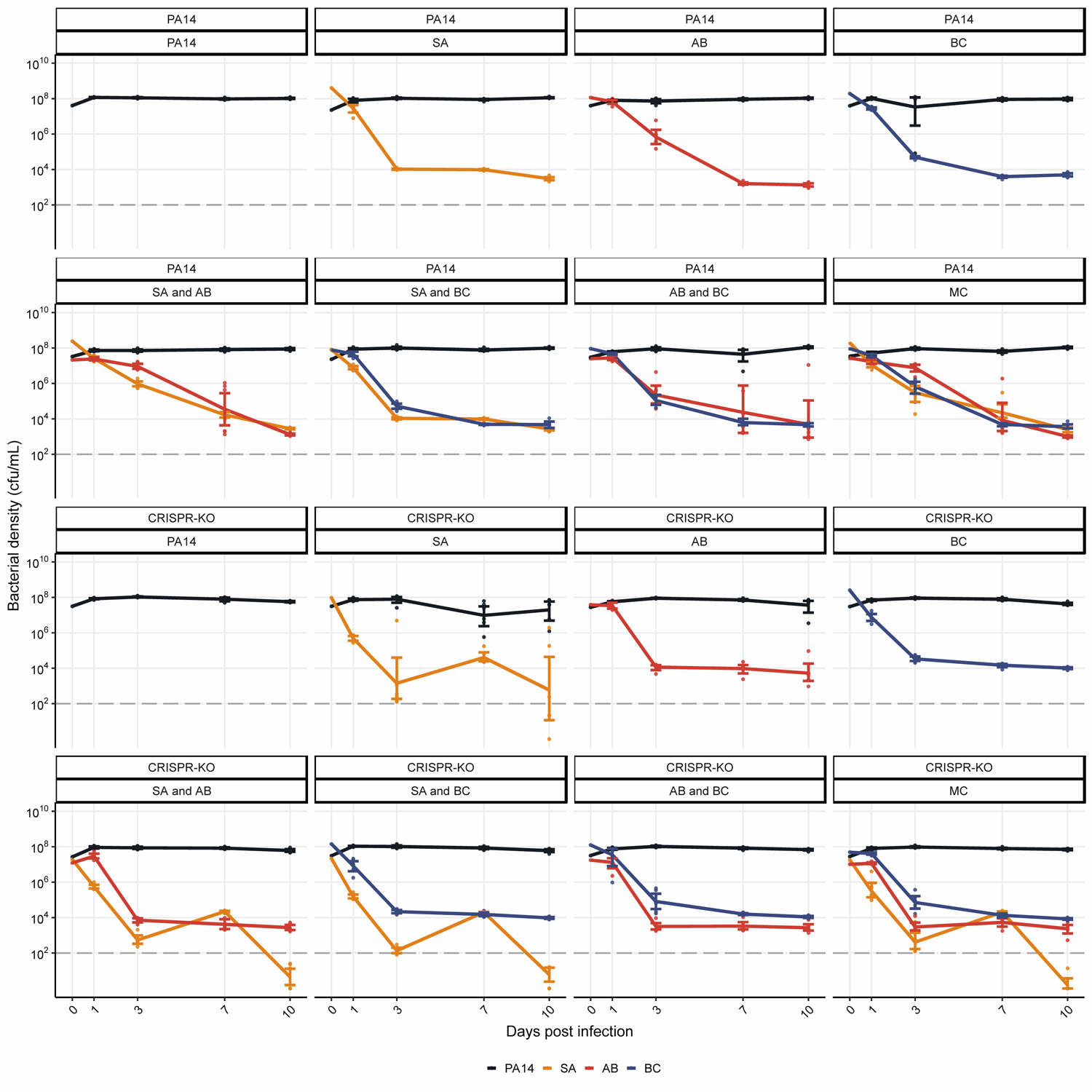
Line plots of bacterial densities in the absence of phage. Showing the bacterial densities in cfu/mL over time for the PA14 WT and CRISPR-KO *P. aeruginosa* strains, and **b** the other microbial community species (SA = *S. aureus*, AB = *A. baumannii*, BC = *B. cenocepacia,* MC = microbial community) in the absence of phage DMS3vir. Dashed horizontal line at 10^2^ cfu/mL marks the threshold of reliable detection where the qPCR results indicate the bacteria has gone or is close to extinction from a population. Data are mean ± 95% CI.

**Supplemental Fig 4.**
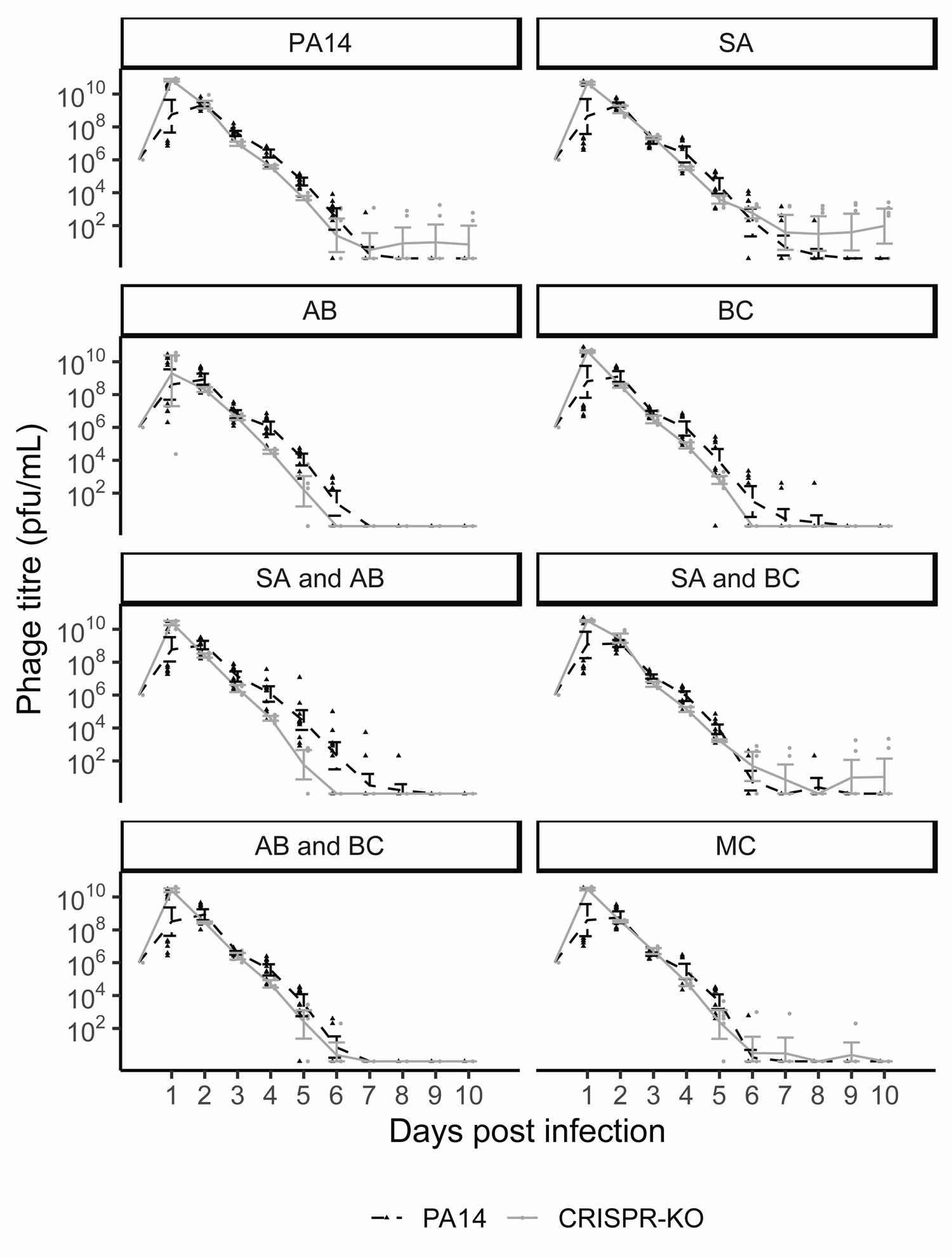
Phage titres over time for each experimental treatment. Phage titres for phage DMS3vir over time across all experimental treatments (SA = *S. aureus*, AB = *A. baumannii*, BC = *B. cenocepacia,* MC = microbial community), infecting either the PA14 WT or the CRISPR-KO strain as indicated by line type. Each data point represents a replicate, with lines following the mean and the error bars denoting 95% CI. Asterisks indicate a significant overall difference in phage density between the PA14 WT (n = 12 per timepoint) or CRISPR-KO clone (n = 6 per timepoint) (effect of *P. aeruginosa* clone; linear models: * p < 0.05).

**Supplemental Fig 5.**
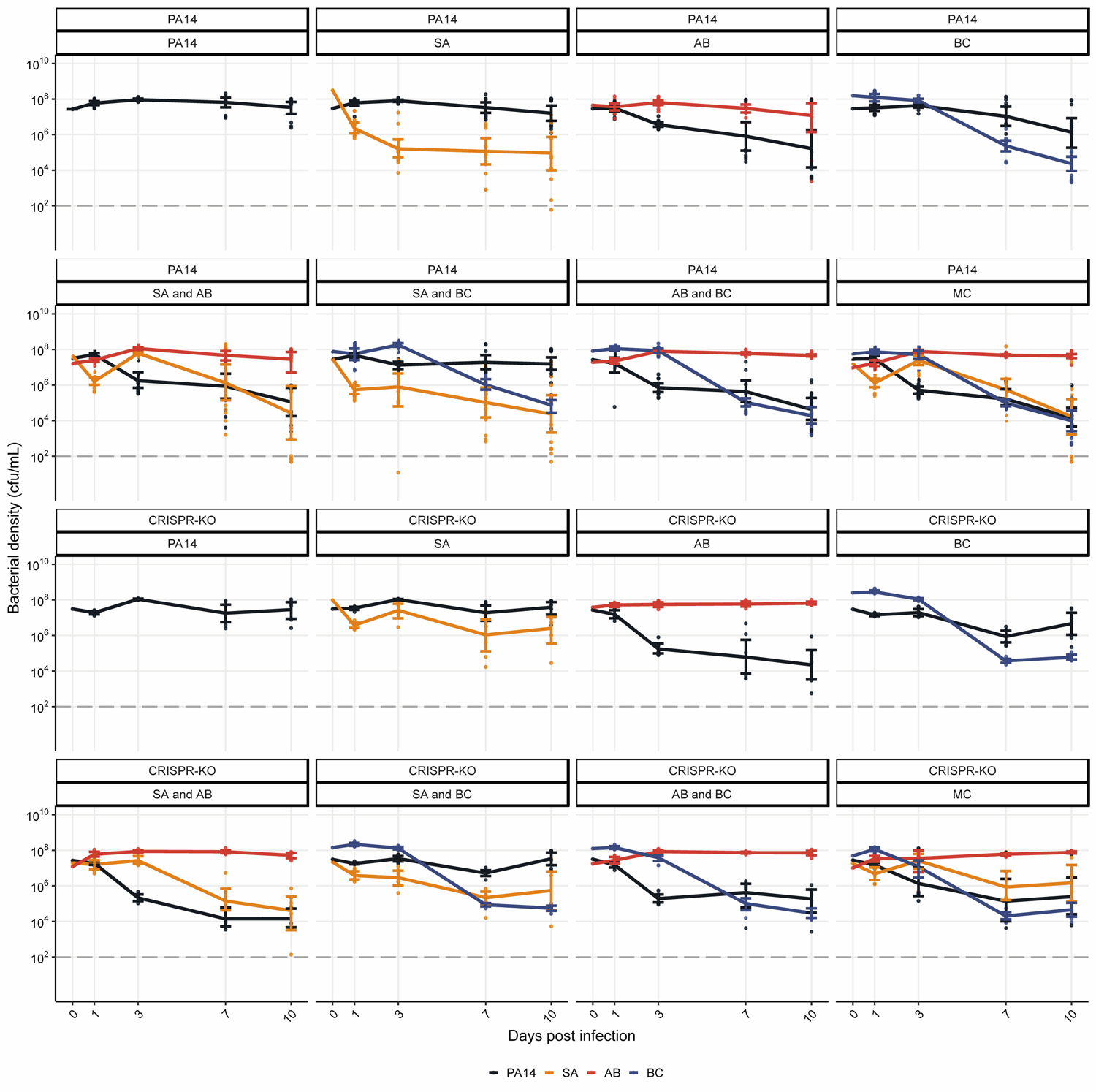
Line plots of bacterial densities in the presence of phage. Showing the bacterial densities in cfu/mL over time for the PA14 WT and CRISPR-KO *P. aeruginosa* strains, and **b** the other microbial community species (SA = *S. aureus*, AB = *A. baumannii*, BC = *B. cenocepacia*, MC = Microbial community) in the presence of phage DMS3vir. Dashed horizontal line at 10^2^ cfu/mL marks the threshold of reliable detection where the qPCR results indicate the bacteria has gone or is close to extinction from a population. Data are mean ± 95% CI.

**Supplemental Fig 6.**
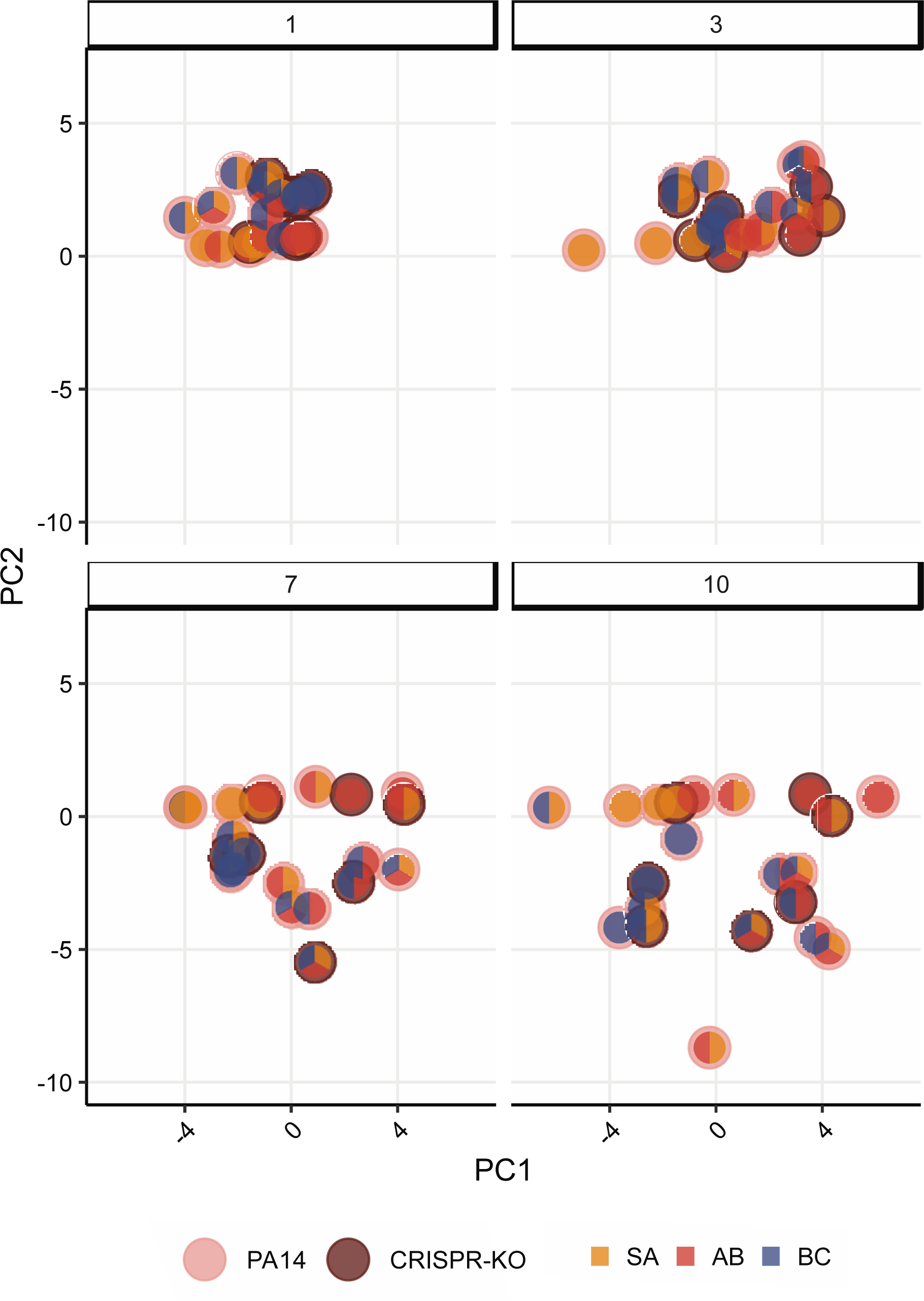
Ordination plots in the presence of phage. PCA ordination of relative bacterial abundance in the presence of phage DMS3vir, with grid layouts separated into days post phage infection. Outer circle colour indicates which PA14 clone is present in the population, while inner circle indicates community composition (SA = *S. aureus*, AB = *A. baumannii*, BC = *B. cenocepacia*).

**Supplemental Fig 7.**
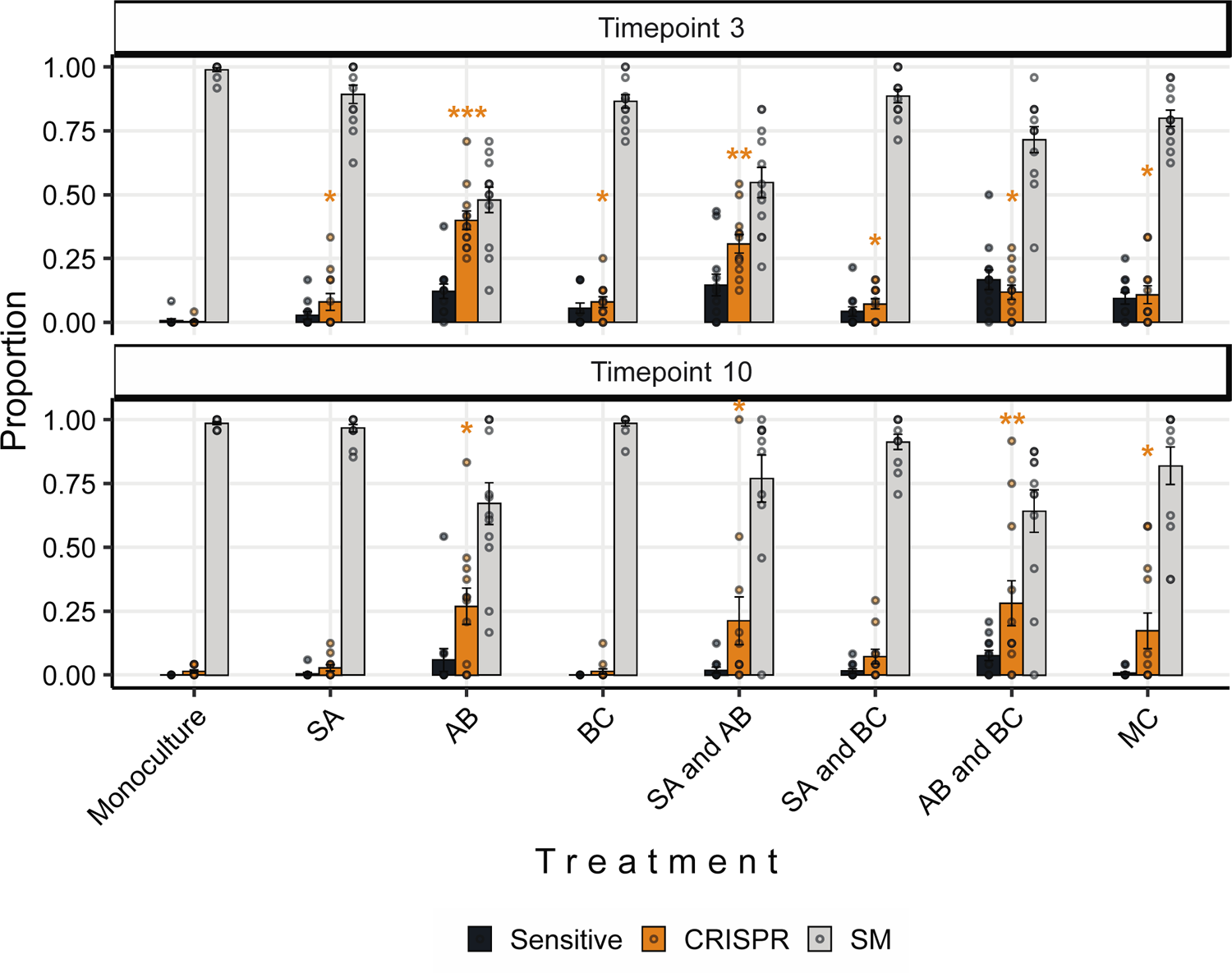
Interspecific competition affects the proportion of evolved CRISPR-based phage resistance. Proportion of *P. aeruginosa* PA14 WT at timepoints 3 and 10 that evolved phage-resistance either through surface modification (SM) or CRISPR immunity, or which remained sensitive to phage DMS3vir when grown in monoculture or different polycultures (SA = *S. aureus*, AB = *A. baumannii*, BC = *B. cenocepacia*). Data are mean ± SE. Asterisks indicate a significant difference in proportion of CRISPR immunity evolved when compared to the PA14 monoculture within each timepoint (n = 12 per treatment) (generalised linear model, quasibinomial: * p < 0.05, ** p < 0.01, *** p < 0.001).

**Supplemental Fig 8.**
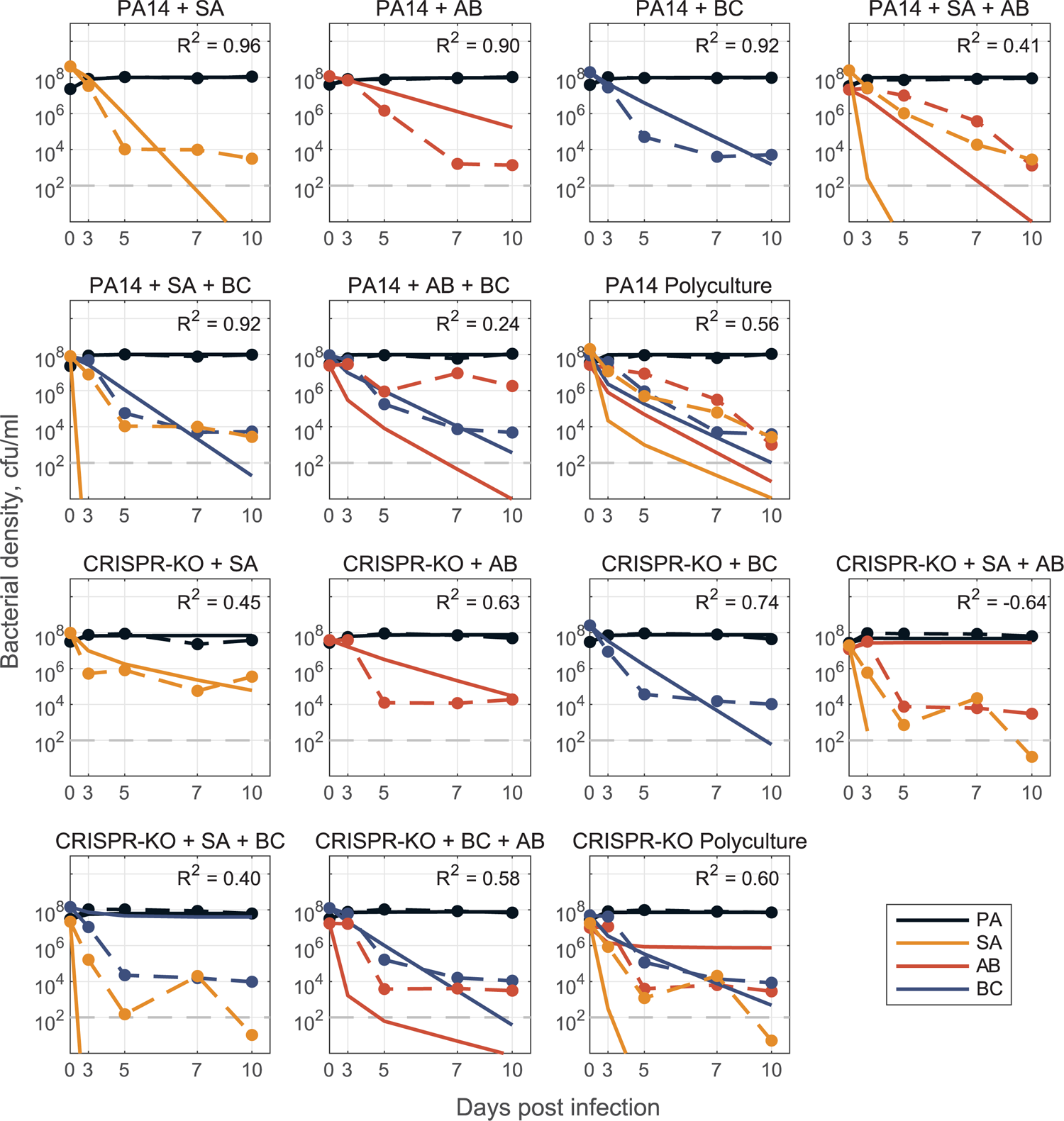
Model from no phage data, trained on only pairwise experimental data. Model fit predictions for two-, three-, and full four species community dynamics (solid lines) compared to experimental data (dashed lines). Models were parameterized via optimization with least-squares to fit a system of ODEs (defined as a generalized Lotka-Volterra competition model with n species, where n=1,2,3,4). We parameterize the models via fitting of 1-(for growth rates *r*_*i*_) and 2-(for all possible pairwise interaction coefficients *β*_*i*,*j*_ ∀*i*,*j* = 1,2) species dynamics and use the resulting coefficients to predict the 3- and 4-species community dynamics. For fitting co-culture data, growth rates *r*_*i*_ were fixed from mono-culture data and interaction parameters *β*_*i*,*j*_ were all open. See Methods and Text S1 for a detailed description of mathematical modelling.

**Supplemental Fig 9.**
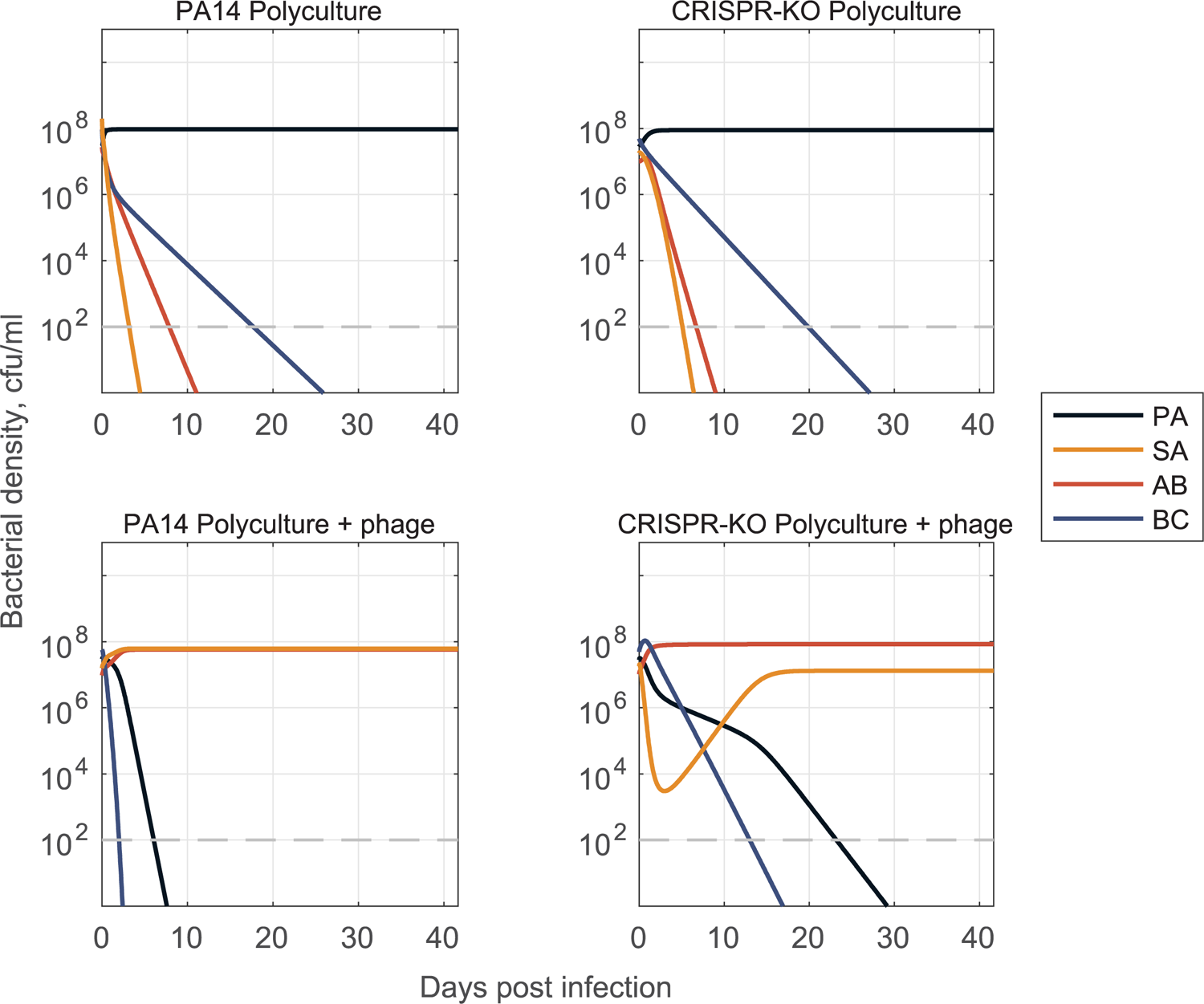
Long time simulation of full community model shows shift in ecological outcomes given inclusion of phage. Simulation of the 4-species community gLV model over a long time scale reveals a qualitative shift in the outcome of the community when phage is present. In the absence of phage (top), *P. aeruginosa* is the dominant competitor and only surviving species. In the presence of phage (bottom), the dominant competitor is eliminated, and we see competitive release of *A. baumannii* and *S. aureus* – maintaining 2 of the 3 non-targeted species in the community. Growth and interaction coefficients for simulation are from the model fits in Figures 7 and 8. For a detailed description of model parameterization and simulation methods, see Methods and Text S1.

